# Changes choroidal area following trabeculectomy: long-term effect of intraocular pressure reduction

**DOI:** 10.1101/486357

**Authors:** Kazuhiro Kojima, Kazuyuki Hirooka, Shozo Sonoda, Taiji Sakamoto, Yoshiaki Kiuchi

**Affiliations:** Department of Ophthalmology and Visual Science, Graduate School of Biomedical Sciences, Hiroshima University, Hiroshima, Japan; Department of Ophthalmology, Kagawa University Faculty of Medicine, Miki, Kagawa, Japan; Department of Ophthalmology, Kagoshima University Graduate School of Medical and Dental Sciences, Kagoshima, Japan

## Abstract

**Purpose:** To investigate the long-term effects of intraocular pressure (IOP) changes after trabeculectomy on the macular and peripapillary choroidal areas.

**Methods:** This prospective longitudinal study examined 30 eyes of 30 patients with glaucoma that was uncontrolled by medical therapy. At 1 day before and at 1 year after the trabeculectomy surgery, macular and peripapillary choroidal images were recorded by enhanced depth imaging optical coherence tomography (EDI-OCT). Luminal and interstitial areas were converted to binary images using the Niblack method. Factors influencing the macular choroidal and peripapillary area were examined by multivariate analysis.

**Results:** After trabeculectomy, the mean IOP was 10.8±3.2 mmHg compared to 17.8±7.2 mmHg at baseline (*P* < 0.001). The total macular choroidal area after the surgery increased from 317,735±77,380 to 338,120±90,700 μm^2^, while the interstitial area increased from 108,598±24,502 to 119,172±31,495 μm^2^ (all *P* < 0.05). The total peripapillary choroidal area after the surgery also increased from 1,557,487±431,798 to 1,650,253±466,672 μm^2^, while the interstitial area increased from 689,891±149,476 to 751,816±162,457 μm^2^ (all *P* < 0.05). However, there were no significant differences observed in the luminal area before and after the surgery. A decrease in the IOP was among the factors associated with the changes in the peripapillary choroidal area.

**Conclusions:** IOP reductions after trabeculectomy led to increases in the macular and peripapillary choroidal areas for at least 1 year postoperative. Increases in the interstitial areas were the primary reason for observed changes in the choroidal area after trabeculectomy.

## Introduction

As metabolic support for the prelaminar portion of the optic nerve head is provided by the choroid,^1–3^ this suggests that it may play an important role in glaucoma.^4–6^ Even though indocyanine green angiography has traditionally been used to visualize choroidal vasculature,^7^ other methods, such as optical coherence tomography (OCT) have also been used to study choroidal morphology. However, potential problems with these previous methods have led to the development of the enhanced depth imaging (EDI) spectral domain OCT method, which makes it possible to perform in vivo cross-sectional imaging of the choroid.^8^

The exact mechanism of glaucomatous optic neuropathy remains unknown, even though glaucoma is one of the leading causes of blindness worldwide. As it has been shown that progression of glaucoma is due to an elevated intraocular pressure (IOP), many studies have demonstrated the benefit of decreasing the IOP.^9,10^ In order to reduce the IOP in glaucoma, trabeculectomy has been used and remains one of the most commonly performed filtration surgeries. Several investigations have reported increases in the subfoveal and peripapillary choroidal thicknesses in primary open-angle glaucoma (POAG) and in primary angle closure glaucoma (PACG) after trabeculectomy-caused IOP reductions.^11–13^ Zhang et al.^14^ recently found that the approximately equal increases in intravascular and extravascular compartments were related to increases in the choroidal thickness that occur after trabeculectomy. Measurements of the choroidal thicknesses in these previous studies were performed at 1.7 mm superior, temporal, inferior, and nasal to the optic disc center and at 1 and 3 mm nasal, temporal, superior, and inferior to the fovea. However, our recent investigation examined a 1,500 μm wide macular choroidal area and a 1.7 mm area around the optic nerve disc center in the peripapillary choroidal area.^15,16^ In contrast to other previous studies, our use of an increased measurement area made it possible to collect greater amounts of information from the choroid. Furthermore, we recently found that the increases that occurred at 2 weeks after trabeculectomy in the macular and peripapillary choroidal areas due to increases in the luminal areas were related to the reduction in the IOP that occurred after the surgery.^15^ However, there is also the possibility that this increase could have been associated with inflammation. To definitively determine the effect of IOP changes on the choroidal area, a long-term follow-up is required. Therefore, the aim of our current study was to investigate the choroidal area changes that occurred at 1 year after the initial trabeculectomy.

## Materials and Methods

### Subjects

Between July 2016 and February 2017, this prospective longitudinal study evaluated the eyes of patients who underwent trabeculectomy treatments at Kagawa University Hospital. Written informed consent was provided by all enrolled subjects in accordance with the principles outlined in the Declaration of Helsinki. The Kagawa University Faculty of Medicine Institutional Review Board approved the study protocol. In addition, prior to patient enrollment, all patients signed the standard consent required for surgery and provided written informed consent to participate in this research study.

This study enrolled all glaucoma patients who had uncontrolled IOP while taking maximally tolerated medication. After enrollment, all subjects underwent visual examinations that included slit lamp, gonioscopy, refraction, and central and peripheral fields. Fornix-based trabeculectomy was performed in all of the patients by one surgeon (KH). Patients enrolled in the study were required to have a spherical refraction within ± 6.0 diopters (D) and a cylinder within ± 2.0 D. Exclusion criteria included having any history of retinal diseases (e.g., diabetic retinopathy, macular degeneration, retinal detachment), having previously undergone laser therapy, exhibited poor image quality due to unstable fixation, or being found to have severe media opacities. Patients having a previous treatment history with medications known to affect retinal thickness (intravitreal anti-VEGF therapy) were also excluded from the study. EDI-OCT examinations in all of the cases were performed by the same investigator.

### EDI-OCT

At 1 day before and 1 year after the surgery, the Heidelberg Spectralis (Heidelberg Engineering, Heidelberg, Germany) with the EDI-OCT technique was used to obtain the macular or peripapillary choroidal images. Measurements were done between 1300-1500 hours. To perform the macular region scans, seven horizontal lines of 30 × 10° through the center of the fovea were used. Scans of the peripapillary region used a 360°, 3.4 mm diameter circle scan that was centered on the optic disc. To obtain each image, an eye tracking system was used, with the best quality image from at least three scans chosen for the subsequent analysis. The area found between the outer portion of the hyperreflective line that corresponded to the retinal pigment epithelium (RPE) and the inner surface of the sclera was defined as the choroidal thickness. In line with the methodology of our previous studies, a masked procedure was used for all of the measurements.^15,16^ All eyes were examined without mydriasis.

### Binarization of the choroid EDI-OCT images

After performing the EDI-OCT evaluation and masking the best images, these images were then displayed on a computer screen. Each of the images were evaluated by one of the authors (HK). Binarization of the choroidal area in each of the EDI-OCT images was performed using the previously described modified Niblack method.^14^ Briefly, the EDI-OCT image was first analyzed using ImageJ software (version 1.47, NIH, Bethesda, MD). For this analysis, we examined a 1,500 μm wide area of the macular choroid that extended vertically to the fovea with 750 μm nasal and 750 μm temporal margins. The ImageJ ROI Manager determined the area to be analyzed, which included a 1.7 mm area that was located around the optic nerve disc center and spanned from the retinal pigment epithelium to the chorioscleral border. After we randomly selected 3 choroidal vessels with lumens > 100 μm through the use the Oval Selection Tool on the ImageJ tool bar, the reflectivities of the lumens were then averaged. The average reflectivity was set as the minimum value in order to reduce the noise in the OCT image. After conversion and adjustment of the image to 8 bits via the use of the Niblack Auto Local Threshold, the binarized image was once again converted to an RGB image. Both the binarization procedures and the automated calculations by the ImageJ software require conversions of the images. Determination of the hyporeflective area was performed using the Threshold Tool, with dark pixels defined as hyporeflective areas, while light pixels were defined as hyperreflective areas. In order to perform the automatic calculations of the hyperreflective and hyporeflective areas, it was necessary to first add data on the relationship between the distance on the fundus and the pitch of the pixels in the EDI-OCT images, which is dependent on the axial length.

### Statistical analysis

All statistical analyses were performed using SPSS for Windows (SPSS Inc., Chicago, IL). A paired *t*-test was used to compare the preoperative and postoperative values. Pearson’s correlation coefficient was used to evaluate the correlation between the choroidal area changes, and the correlations among the choroidal area, systolic blood pressure (SBP), diastolic blood pressure (DBP), IOP, age, and axial length. A subsequent multivariate regression analysis was performed using variables that had a Pearson’s correlation coefficient value of *P* < 0.2. The choroidal area was defined as the dependent parameter for the multivariate analysis, while the independent parameters included the other parameters selected by the Pearson’s correlation coefficient and the choroidal area. *P* < 0.05 was considered statistically significant. All statistical values are presented as the mean ± standard deviation (SD).

## Results

Table 1 shows the clinical characteristics of the 30 eyes of 30 patients enrolled in the study. The mean age of the patients was 68.9±9.0 years (range: 50 to 87 years).

**Table 1.**
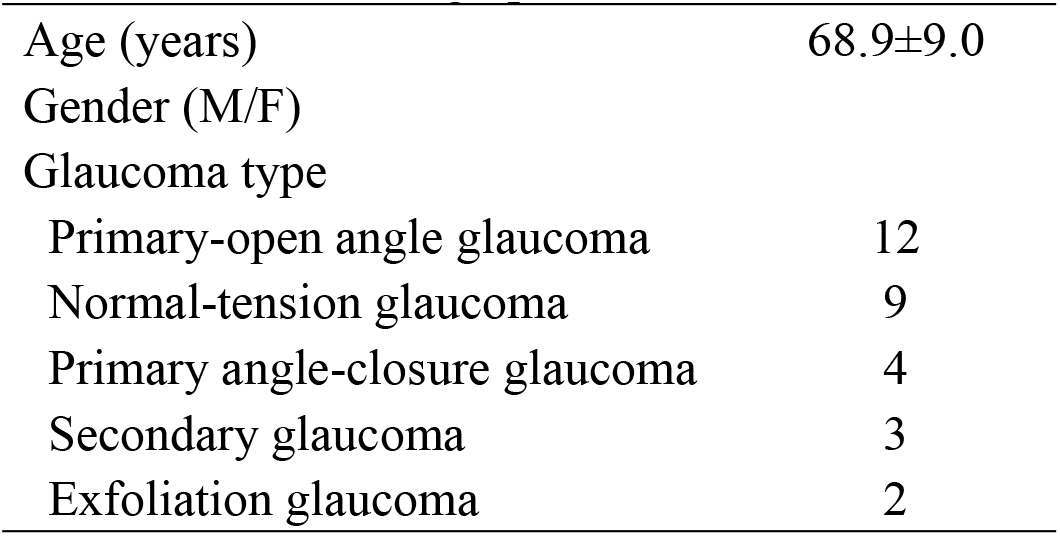
Patient demographic and clinical data

|  |  |
| --- | --- |
| Age (years) | $68.9 \pm 9.0$ |
| Gender (M/F) |  |
| Glaucoma type |  |
| Primary-open angle glaucoma | 12 |
| Normal-tension glaucoma | 9 |
| Primary angle-closure glaucoma | 4 |
| Secondary glaucoma | 3 |
| Exfoliation glaucoma | 2 |

The mean IOP decreased from 17.8±7.2 to 10.8±3.2 mmHg (*P* < 0.001), while the mean OPP increased from 45.2±11.3 to 57.0±9.2 mmHg (*P* < 0.001; Table 2) after the trabeculectomy. After the surgery, the axial length decreased from 24.5±1.5 mm before surgery to 24.2±1.4 mm after the surgery (*P* < 0.001; Table 2).

**Table 2.** IOP, BP, OPP and axial length before and after trabeculectomy

|  | Before | After | *P* value |
| --- | --- | --- | --- |
| IOP (mmHg) | $17.8 \pm 7.2$ | $10.8 \pm 3.2$ | $<0.01$ |
| BP (mmHg) |  |  |  |
| Systolic | $127.5 \pm 19.8$ | $138.2 \pm 17.8$ | 0.02 |
| Diastolic | $78.0 \pm 13.9$ | $84.5 \pm 13.2$ | 0.07 |
| OPP (mmHg) | $45.2 \pm 11.3$ | $57.0 \pm 9.2$ | $<0.01$ |
| Axial length (mm) | $24.5 \pm 1.5$ | $24.2 \pm 1.4$ | $<0.01$ |

IOP; intraocular pressure, BP; blood pressure, OPP; ocular perfusion pressure

After the surgery, the macular choroidal area increased, with the total area increasing from 317,735±77,380 to 338,120±90,700 μm^2^, while the interstitial area increased from 108,598±24,502 to 119,172±31,495 μm^2^ (all *P* < 0.05, Table 3). The peripapillary choroidal area also exhibited increases after the surgery, with the total area increasing from 1,557,487±431,798 to 1,650,253±466,672 μm^2^, while the interstitial area increased from 689,891±149,476 to 751,816±162,457 μm^2^ (all *P* < 0.05). However, no significant differences were noted for the luminal area before and after the surgery.

**Table 3.** Choroidal area observed on EDI-OCT images before and after surgery

|  | Macula |  |  | Peripapilla |  |  |
| --- | --- | --- | --- | --- | --- | --- |
|  | Before | After | *P* value | Before | After | *P* value |
| Total area ($\mu\text{m}^2$) | 317,735±77,380 | 338,120±90,700 | 0.03 | 1,557,487±431,798 | 1,650,253±466,672 | 0.03 |
| Luminal area ($\mu\text{m}^2$) | 209,137±56,767 | 218,948±61,424 | 0.15 | 867,596±301,209 | 898,437±312,174 | 0.28 |
| Interstitial area ($\mu\text{m}^2$) | 108,598±24,502 | 119,172±31,495 | 0.01 | 689,891±149,476 | 751,816±162,457 | 0.001 |

A positive correlation was observed for the magnitude of change between the macular choroidal area and the OPP (r = 0.44, *P* = 0.02; Table 4). However, there was no observed correlation for the magnitude of the change between the macular choroidal area and the IOP reduction (r = −0.32, *P* = 0.09). In contrast, a negative correlation was observed for the magnitude of the change between the peripapillary choroidal area and the IOP reduction (r = −0.58, *P* < 0.01; Table 5).

**Table 4.** Pearson's correlation between changes in the magnitude for the macular choroidal area and each factor

|  | r | P value |
| --- | --- | --- |
| Age | -0.27 | 0.89 |
| Changes in SBP | 0.34 | 0.08 |
| Changes in DBP | 0.20 | 0.30 |
| Changes in IOP | -0.32 | 0.09 |
| Changes in AL | -0.33 | 0.11 |

SBP: Systolic blood pressure, DBP: Diastolic blood pressure, IOP: Intraocular pressure, AL: Axial length

**Table 5.** Pearson's correlation between the magnitude change for the peripapillary choroidal area and each factor

|  | r | P value |
| --- | --- | --- |
| Age | 0.12 | 0.52 |
| Changes in SBP | 0.26 | 0.19 |
| Changes in DBP | 0.03 | 0.87 |
| Changes in IOP | -0.58 | <0.01 |
| Changes in AL | -0.40 | 0.05 |

SBP: Systolic blood pressure, DBP: Diastolic blood pressure, IOP: Intraocular pressure, AL: Axial length

Factors that could potentially influence the increases observed in the macular or peripapillary choroidal area were also investigated. Table 6 presents the results of the multivariate analyses for each parameter. These findings showed that there were no significant correlations observed for the changes in the macular choroidal area. However, our analyses did find that there was a significant association between the changes in the IOP and those in the peripapillary choroidal area (Table 7).

**Table 6.**
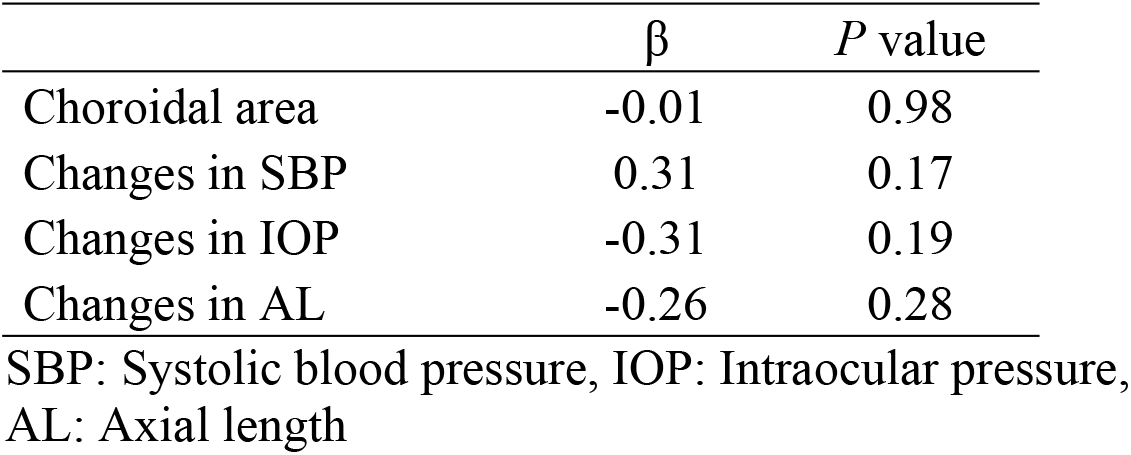
Multivariate analysis of the changes in the association in the subfoveal choroidal area and each factor

**Table 7.** Multivariate analysis of the changes in the Associations in the peripapillary choroidal area and each factor

| | $\beta$ | P value |
| --- | --- | --- |
| Choroidal area | -0.01 | 0.99 |
| Changes in SBP | 0.23 | 0.23 |
| Changes in IOP | -0.53 | 0.02 |
| Changes in AL | -0.19 | 0.36 |

SBP: Systolic blood pressure, IOP: Intraocular pressure, AL: Axial length

## Discussion

Our current study demonstrated that there were increases after trabeculectomy in the long-term subfoveal and peripapillary choroidal areas, with the trabeculectomy also leading to decreases in the IOP. In addition, we also determined that there were increases in both the macular and peripapillary choroidal areas, which led to an increase in the interstitium of the choroid.

Kara et al.^11^ reported that at 1 month after trabeculectomy there was a large decrease in the IOP that subsequently led to choroidal thickening. Kadziauskiene et al.^13^ additionally found that the increase in the subfoveal and peripapillary choroidal thickness that occurred after the trabeculectomy for at least 6 months postoperatively was correlated with greater IOP reduction and axial length shortening. Furthermore, we recently reported that the increases in the macular and peripapillary choroidal areas that were caused by the reduction in the IOP at 2 weeks after trabeculectomy were correlated with the subsequent changes in the IOP.^15^ In contrast, although there were short-term (7 days) increases in the choroidal thickness following trabeculectomy in PACG, these changes were found not to be related to either a decrease in the IOP or shortened axial length.^12^ Usui et al.^17^ additionally reported that while the choroid was thicker, the axial length was shorter, and the IOP was lower at 6 days after trabeculectomy, there was no correlation in POAG patients for the IOP changes and the changes in the choroidal thickness at the subfovea. Moreover, during the early stages following trabeculectomy, there was no significant change in the choroidal thickness in accordance with the decreasing IOP. However, for at least 1 year after the trabeculectomy, the increase in the choroidal area (thickness) that occurred after a large decrease in the IOP was correlated with the IOP reduction during the late stages.

We previously reported that increases in the luminal areas that led to increases in the macular and peripapillary choroidal areas were related to reductions in the IOP that occurred at 2 weeks after trabeculectomy.^15^ In the current study, however, increases in the interstitial areas that led to increases in the macular and peripapillary choroidal areas were due to a reduction in the IOP at 1 year after the initial trabeculectomy. Furthermore, our previous study showed that the rate of increase in the macular choroidal interstitial area or luminal area at 2 weeks after trabeculectomy was 111.2% or 118.6%, respectively, while the rate of increase for the peripapillary choroidal interstitial area or luminal area was 112.0% or 128.2%, respectively.^15^ In contrast, the rate of increase in the macular choroidal interstitial area or luminal area at 1 year after trabeculectomy was 109.1% or 104.7%, respectively, while the rate of increase for the peripapillary choroidal interstitial area or luminal area was 109.0% or 103.5%, respectively. Zhang et al.^14^ additionally reported that at 6 months after the trabeculectomy, choroidal thickness increases were observed in conjunction with decreasing IOP in both the large choroidal vessels and interstitium of the choroid. Furthermore, the luminal area returned to the originally observed size seen prior to the surgery even though the IOP reduction at 1 year after the trabeculectomy caused an increased choroidal area. Another recent study that examined choroidal vessels, also reported finding that the luminal area changed in accordance with the diurnal variation.^18^ Thus, this suggests that changes could easily occur in the luminal area. Moreover, other studies have reported that the ocular blood flow increases seen after trabeculectomy can also potentially contribute to thickening of the choroid.^11,19^ Therefore, the question that needs to be answered is, do increases in the ocular blood flow still occur even after a return of the luminal area to its pre-surgery size? However, since we did not measure the ocular blood flow, it was not possible to determine this in our current study. Further studies that examine the blood supply of the optic nerve disc after IOP reduction following trabeculectomy, especially in the peripapillary area, will need to be undertaken.

There were several limitations for our current study. First, this study only examined a small number of subjects. Thus, a further study with a larger number of patients will need to be undertaken in order to address this issue. Second, since manual segmentation cannot achieve perfect reproducibility, a truly objective method would be preferable in this type of study.

## Conclusions

The present study demonstrated that IOP reduction after trabeculectomy led to an increase in the macular and peripapillary choroidal areas, with these increases continuing for at least 1 year. These noted increases were found to be due to an increase in the interstitial areas. Thus, overall our findings demonstrated that changes in the IOP were significantly associated with changes in the peripapillary choroidal area.

## Acknowledgements

The authors thank FORTE for the professional service that edited our manuscript.

